# memVar: an integrated resource for interpreting human membrane protein variation in molecular and disease contexts

**DOI:** 10.64898/2026.11.04.764121

**Authors:** Yizhou Xu, Siming Huang, Jingyao Zhao, Jianfeng Sun

## Abstract

Interpreting membrane protein variation requires connecting clinical evidence to the membrane environments and molecular interactions that govern protein functions. Here, we introduce memVar (https://memvar.sunnylab.org), an integrated database of 10,906,688 unique single-nucleotide variants from ClinVar, COSMIC, and gnomAD across 7,732 human membrane-associated proteins. memVar reconciles genomic, transcript, and protein coordinates to link variants with membrane topology, bilayer depth, lipid contacts, functional sites, partner-specific interaction interfaces, and variant-effect predictions, alongside disease, tissue-expression and regulatory annotations. Coordinated sequence, structure and variant views retain source provenance and distinguish clinical assertions, experimental observations, and computational predictions. Our analysis reveals that pathogenic/likely pathogenic missense substitutions are enriched in transmembrane segments and predicted protein-protein interaction sites relative to benign/likely benign substitutions, with pooled odds ratios of 2.47 and 6.67, respectively. IntAct-derived interaction loss or decrease was experimentally recorded for 57.9% of pathogenic/likely pathogenic substitutions versus 7.4% of benign/likely benign substitutions. CFTR examples contextualize the established G551D gating defect and prioritize W202R, a variant of uncertain significance, for tests of protein maturation, surface abundance and channel activity. By connecting variant records to specific molecular questions, memVar potentially supports mechanism-oriented investigation and prioritization of functional experiments.

**Graphical abstract:** 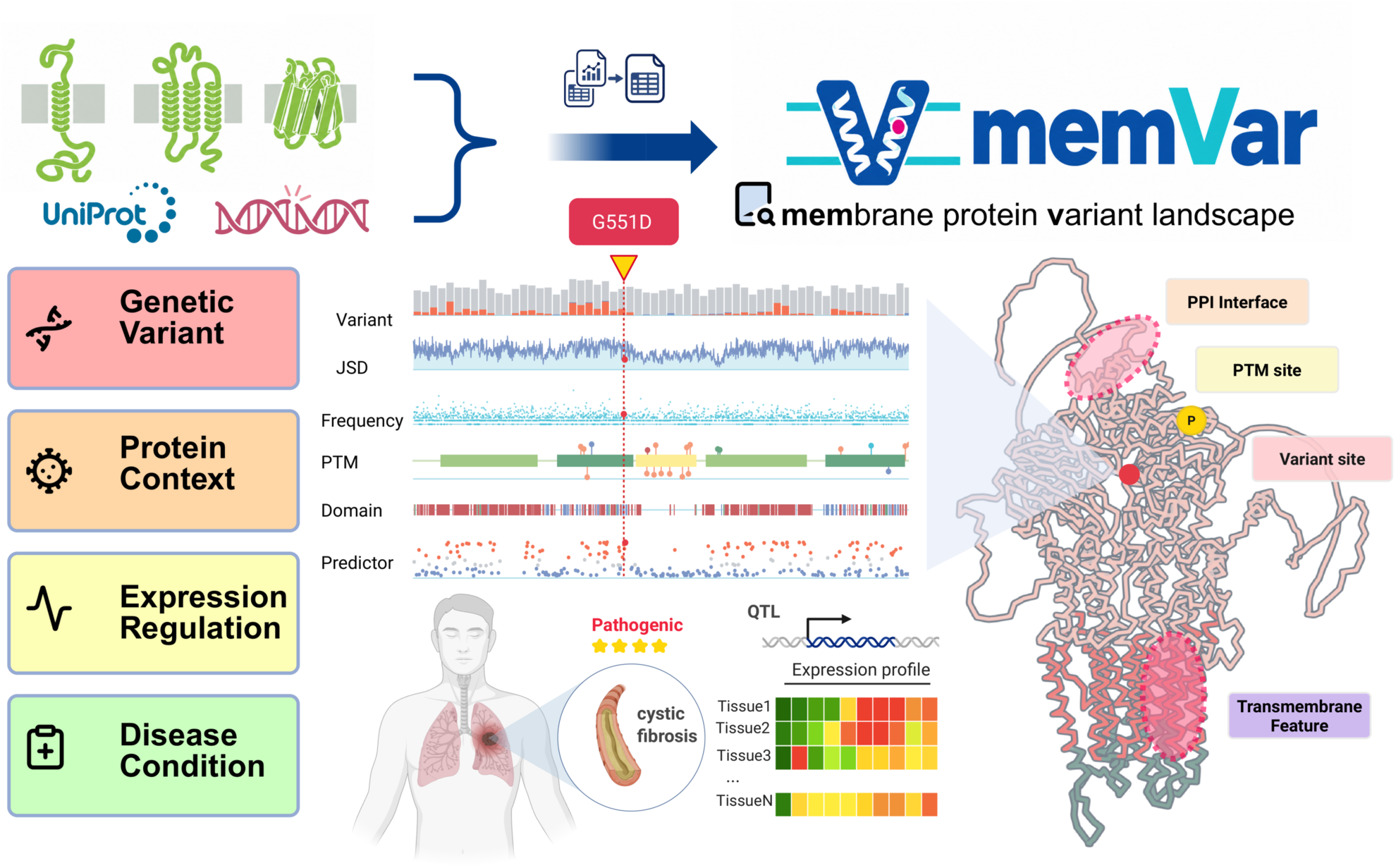

## Introduction

Decades of advances in structural biology have expanded the Protein Data Bank (PDB) (1) to an archive of more than 250,000 experimentally determined macromolecular structures, a milestone reached in March 2026 (2). Meanwhile, our knowledge of human variation has grown substantially, manifested by several trustworthy databases, such as ClinVar, first fully released in 2013, with more than 4.5 million variant records (3), and gnomAD v4 aggregating variant cohorts based on sequencing data from 807,162 individuals (4). Despite these advances, the structural information required to interpret genetic variation remains distributed across protein classes. A recent assessment estimated that transmembrane proteins account for only approximately 5% of high-resolution experimental structures in the PDB (5). Thus, although clinical and population databases document extensive membrane protein variants, understanding its molecular consequences requires integrating genetic evidence with membrane topology, residue-level features, and functionally relevant interaction interfaces.

Approximately one quarter of human protein-coding genes encode membrane proteins, which mediate ion homeostasis, nutrient transport and cellular communication (6, 7). Their pharmaceutical importance has long been established in the prominence of G protein-coupled receptors (GPCRs). For instance, a recent study identified 516 approved drugs acting at 121 GPCRs, representing 36% of approved drugs in that survey (8), while GPCR-directed therapies for diabetes and obesity illustrate the commercial value of this target class (9). Understanding variant effects in membrane proteins can also inform therapeutic intervention. For example, the pathogenic CFTR G551D variant impairs channel gating. This defect illuminates the rationale for pharmacological potentiation, which clinical studies subsequently evaluated in patients carrying that variant (10, 11). Mechanistic insight into variant effects therefore informs disease interpretation and guides therapeutic target modulation.

The lipid bilayer adds a distinct physical constraint to membrane protein variation, i.e., the energetic consequences of a substitution depend on the position of a residue within the membrane (12–14). Variants can therefore disrupt membrane insertion, topology, helix packing or lipid contacts, with consequences for trafficking, gating, transport and signalling (15–17). Pathogenic substitutions also occur outside the bilayer, as exemplified by CFTR G551D in a cytoplasmic nucleotide-binding domain (10). Interpretation thus requires full-length protein coverage coupled to residue-specific membrane context. Existing resources, such as TMFunction (18), MutHTP (19), TMSNP (20), memPmut (21), and MPTherm (22), catalogue membrane protein mutations and phenotypes, experimental stability and binding measurements (22, 23), membrane orientation and topology (24, 25), or protein and clinical annotations (26, 27).

Clinical and experimental evidence covers only a small fraction of possible human missense variants, which creates a need for computational approaches that predict variant effects at scale. Recent progress in variant-effect prediction makes the integration increasingly consequential. A workflow based on the ESM1b protein language model predicted the effects of approximately 450 million possible missense variants across 42,336 human protein isoforms (28), while AlphaMissense supplied predictions for 71 million missense variants accessible through single-nucleotide changes (29). These advances have extended variant assessment far beyond current experimental coverage. For membrane proteins, an independent evaluation of AlphaMissense highlighted the importance of reference annotation quality when assessing prediction performance (30). Dedicated approaches, including Pred-MutHTP (31), Missense3D-TM (32), MutDPAL (33), and Memo-Patho (34), have further expanded the available methods for evaluating variants in membrane environments. However, experimental findings, clinical annotations and these increasingly diverse predictions remain distributed across resources, limiting their joint interpretation within the relevant molecular context. This gap motivates a unified membrane-focused resource that brings these complementary data together with membrane topology, protein structure and functional annotations. Moreover, molecular interaction networks, gene expression levels, and regulatory annotations illuminate the biological context of genetic variation (35–38). The central challenge lies in connecting this broader context to the mechanisms through which individual variants alter protein function.

Here, we present memVar, a database designed for the systematic investigation of variation in the human membrane proteome. memVar brings together 10,906,688 unique single-nucleotide variants across 7,732 proteins, including transmembrane, lipid-anchored, and peripheral membrane proteins. It maps variants across genomic, transcript, and protein coordinates and connects them to membrane features, residue annotations, interfacial profiles, and prediction scores. We also incorporate tissue expression profiles, quantitative trait locus (QTL) associations, and disease annotations to provide biological context. We characterize the resulting catalogue, examine associations between clinical classifications and molecular features, and use CFTR as a case study to demonstrate how integrated evidence can inform testable hypotheses for functional investigation. Together, memVar provides an extensible framework for systematically assessing membrane protein variation in their sequence and biological contexts to guide and prioritize unresolved mechanisms for future curation and experimental validation.

## Materials and methods

### Reference proteins and source integration

We assembled the reference collection from reviewed human reference-proteome (UP000005640) entries carrying the Membrane keyword (KW-0472) in UniProtKB/Swiss-Prot 2026_03 (39). The resulting set comprised 7,732 protein accessions linked to 7,707 HGNC identifiers. Canonical and non-canonical sequences were retained with their UniProt identifiers, giving a total of 16,672 sequences. Gene and transcript relationships were extracted from HGNC, Ensembl and RefSeq cross-references based on Uniprot.

Proteins were assigned mutually exclusive membrane classes based on UniProt annotations and keywords, i.e., integral, peripheral, lipid-anchored, and attachment mechanism unannotated. External records were linked through explicit gene or protein identifiers. Residue-level projection additionally required compatible sequence and positional information, while unresolved mappings were retained as source records without assignment to a canonical residue. Source identity, evidence type and missingness were preserved throughout integration (see **Supplementary Tables S1-S2** for incorporated resources, versions and coverage).

### Variant processing and annotation

We normalized ClinVar (3), COSMIC (40), and gnomAD exome SNV records against GRCh38 using bcftools (41). Unique SNVs were identified by chromosome, position, reference allele, and alternate allele, with source labels and annotations retained. Specific records were selected through transcript or UniProt accession cross-references and reannotated using Ensembl Variant Effect Predictor (VEP) 116 (42) while MANE 1.5 provided RefSeq-Ensembl transcript correspondences (43). We then verified the exact full-length sequence matching against UniProt canonical and isoform sequences, with additional checks of positions and reference residues in HGVS protein descriptions (HGVSp) where applicable. Each protein-variant entry was defined by a UniProt sequence identifier, an SNV, and the full HGVSp expression with its original source annotation recorded. As a result, memVar comprised 10,906,688 unique SNVs represented by 18,892,222 protein-variant entries, documenting 12,782,398 missense (67.66%), 5,441,814 synonymous (28.80%) and 599,034 stop-gained (3.17%) entries.

Population allele frequencies from gnomAD v4.1.1 exomes, Single Nucleotide Polymorphism Database (dbSNP) (44) and precomputed AlphaGenome Atlas variant-impact and splicing scores were assigned by exact genomic allele matching on GRCh38 (45). dbNSFP 5.4a supplied *In silico* predictions of variant-effect scores, conservation metrics, and separate gene-constraint annotations from 18 selected predictors (46, 47). Transcript-dependent scores additionally required agreement with the selected transcript and amino acid substitution, while unresolved values were not replaced by scores from another isoform. The coverage determined by effect predictors was distinguished from the coverage determined by background conservation metrics (see **Supplementary Methods S2** for score-matching and missing-value rules).

### Sequence, structural, and interaction annotations

Membrane topology, orientation, contacts, functional sites, and post-translational modifications collected from Uniprot and specialized databases were mapped to reference sequences using source coordinates, verified sequence context or structure-to-sequence correspondences. Records with ambiguous positions were excluded from analyses requiring exact residue assignments. HMMER identified Pfam domains (48), while DeepTMHMM2 predicted transmembrane (TM) segments, re-entrant (RE) regions and interfacial (IF) helices for all collected sequences (49). Jensen-Shannon divergence (JSD) and Zvelebil scores were calculated from UniRef90 homolog alignments generated with MMseqs2 (50) and filtered with hhfilter. AlphaFold DB structures were checked against their reference sequences (51). ThermoMPNN estimated substitution-associated stability changes only where canonical sequence, structure and residue identity agreed (52). Detailed mapping, alignment and fragment-selection procedures are given in **Supplementary Methods S3**.

BioGRID (53) and IntAct (54) interactions were linked to reference through proteins identifiers while retaining assay, experimental-system and source annotations. SPPIDER-seq predicted partner-specific residue interfaces for eligible canonical sequence pairs (55), and PeSTo supplied structure-based binding propensities (56). International Molecular Exchange (IMEx) mutation-effect records distributed through IntAct were stored separately from ordinary interaction pairs and underwent additional sequence verification before substitution-level comparisons (57). Protein and gene annotations supplied functional, pathway, localization and pharmacological context (**Supplementary Methods S4**).

### Disease, tissue and regulatory data integration

To provide a consistent disease and anatomical context for variant effect interpretation, we linked disease and phenotype annotations through explicit gene and disease identifiers. To accomplish this process, two resources were used, Mondo disease ontology (MONDO) was used to unify disease categories (58), and Uberon multispecies anatomy ontology (UBERON) was used to provide the anatomical 15 groups and for compatible links to expression tissues (59). Multiple disease-group memberships and unresolved mappings were retained (**Supplementary Methods S5**). Expression data from HPA, GTEx and PaxDb were linked through gene or protein identifiers, with tissue labels, measurement types and units retained. RNA expression and protein abundance remained separate measurements. QTL records were associated with their target genes while retaining tissue, study, association statistics and reference assembly. Gene linkage alone did not establish an exact variant match.

### Statistical analysis

We conducted a variety of statistical analyses using pathogenic/likely pathogenic (P/LP) and benign/likely benign (B/LB) missense protein-variant entries in transmembrane proteins. Entries with conflicting or mixed labels were excluded from P/LP-versus-B/LB comparisons. We applied no additional filters based on ClinVar review status or origin filter.

Molecular comparisons were made by using unique, sequence-verified amino acid substitutions within annotation-specific evaluable sets. We compared P/LP and B/LB substitutions for overlap with UniProt transmembrane segments, DeepTMHMM2 membrane regions, and SPPIDER-seq predicted interfaces, as well as for recorded IntAct interaction effects. Conditional exact odds ratios (ORs) were estimated with 95% confidence intervals (CIs), and two-sided Fisher’s exact tests were applied. Bonferroni correction was applied separately to the three primary regional comparisons and the four DeepTMHMM2 categories. We also compared evolutionary conservation and population allele frequencies between the two groups. Conservation comparisons used JSD scores matched to UniProt sequences and residue positions. Population-frequency distributions included entries with positive gnomAD overall allele frequencies.

We evaluated six dbNSFP-derived scores, AlphaGenome Atlas AVI and ThermoMPNN against ClinVar P/LP and B/LB classifications using a common set of protein-variant entries with all eight scores available. Discrimination was assessed by the area under the receiver operating characteristic curve (AUROC) and average precision. For average precision, 95% CIs were estimated from 1,000 paired bootstrap replicates resampling protein accessions. Detailed feature definitions, exclusions, and statistical procedures are provided in the **Supplementary Material**.

### Structural support for variant interpretation

For each protein sequence, memVar displays experimentally determined structures where available and predicted models otherwise. In addition to AlphaFold-predicted structures (60) we also supplied 3D models predicted by ColabFold to broaden the available structural predictions and support comparative assessment of model consistency and confidence (61). Together with sequence, membrane and functional annotations, these structures provide a one-stop resource for examining variant sites and prioritizing functional experiments, reducing the need for independent structure prediction and searches across multiple databases.

### Database implementation

Python pipelines generated Parquet entity and relationship tables, queried through DuckDB and FastAPI. The React, TypeScript and Vite interface uses Nightingale and PDBe Mol* to connect sequence and structural views. AlphaGenome reference tracks are stored in HDF5 and retrieved by genomic interval. Shared protein, gene, variant and coordinate identifiers connect the annotation layers while retaining provenance (**Supplementary Methods S8**).

## Results

### A common reference for membrane protein variation

Overall, memVar systematically integrates genomic variation from ClinVar, COSMIC and gnomAD across human membrane proteins, encompassing single-pass and multi-pass transmembrane proteins as well as lipid-anchored and peripheral membrane proteins (**Figure 1**). By extending beyond proteins that traverse the lipid bilayer, the resource brings together clinical classifications, cancer-associated observations and population allele frequencies within a unified framework, which enabling complementary evidence associated with individual genomic alleles to be examined collectively.

**Figure 1.**
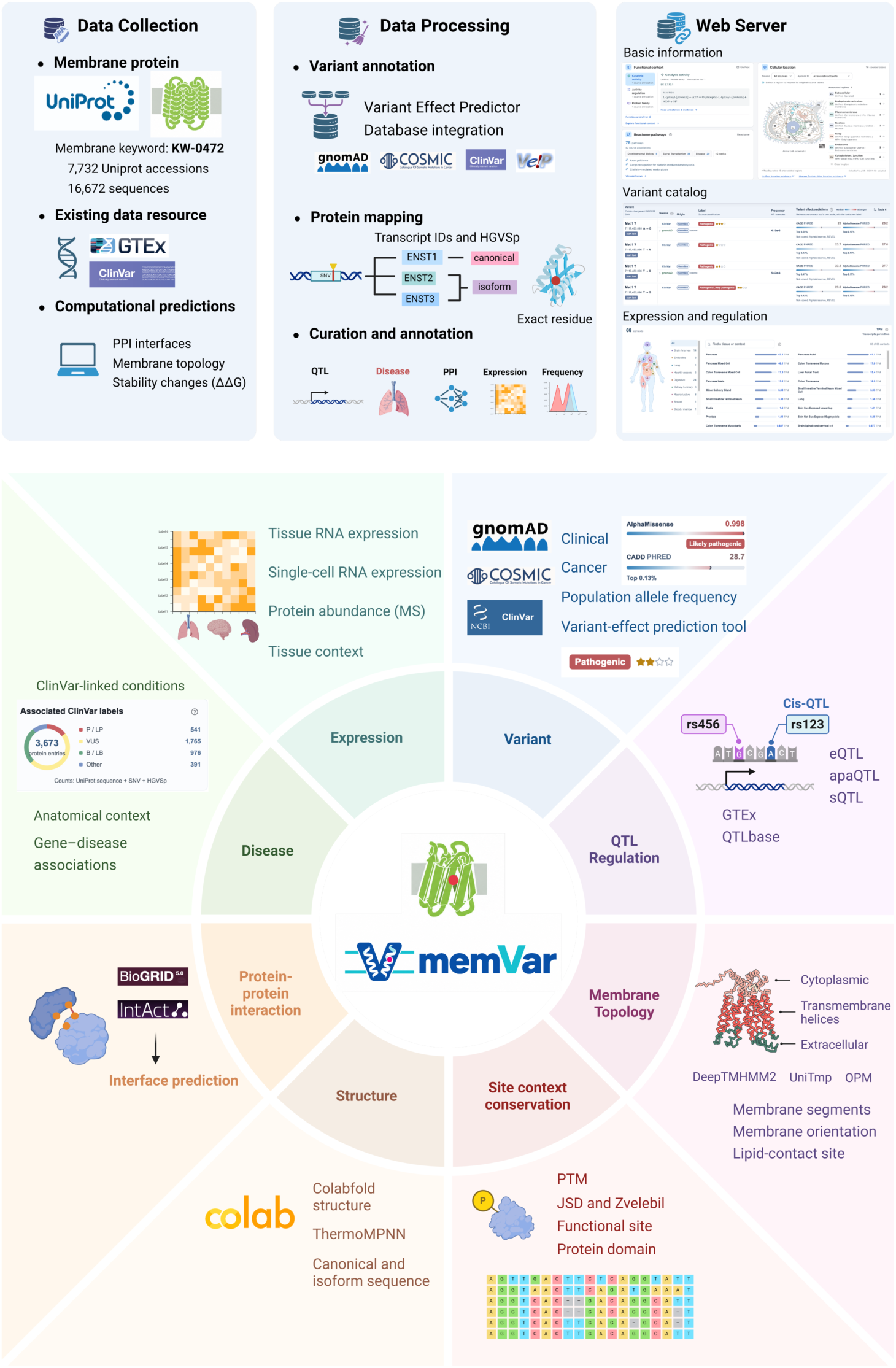
Overview of memVar data integration and exploration. Membrane-associated proteins and isoforms are collected from UniProt, and variants from ClinVar, COSMIC and gnomAD are mapped through transcripts to verified protein sequences. The upper panels summarize data collection, processing and web access. The lower diagram shows the connected annotation layers, including membrane features, structures, functional sites, post-translational modifications (PTMs), conservation, protein-protein interactions (PPIs), disease, expression, and quantitative trait loci (QTLs). JSD, Jensen-Shannon divergence. MS, mass spectrometry.

In memVar, we systematically added sequence and structurally contextual information to residues that possess clinically and experimentally observed mutations. This includes membrane topology, orientation, and lipid contacts, which together define the local environment of a membrane protein position of interest, in which an amino acid substitution occurs. Evolutionary conservation, functional domains, interaction sites, and post-translational modifications are either collected from trustworthy databases/literature or predicted by widely used tools. Native/predicted atomic-level structures, coupled with stability changes in them, collectively support inspection of local packing, while interfacial information connect the residue to possible molecular partners. These are strengthened by a systematic collection of associated functional, pathway, and pharmacological information.

Although the annotations above identify molecular features potentially affected by a variant, they do not establish the tissue or disease contexts in which these effects may be relevant. To bridge this gap, memVar integrates gene-disease associations, tissue-expression profiles, and QTL records, and connects them with broader biological and clinical contexts. Importantly, the integration preserves the original data sources, evidence levels, and coordinate-mapping status, enabling users to trace variant-to-protein relationships and assess the evidence supporting their potential functional and clinical implications.

### Variant composition and annotation coverage

To characterize the breadth of memVar and the distribution of evidence available for variant interpretation, we systematically examined the overlap among variant repositories, functional consequences across canonical and isoform sequences, membrane-protein classifications, and coverage of integrated molecular annotations (**Figure 2**). Integration of gnomAD, ClinVar and COSMIC yielded 10,906,688 unique genomic SNVs. Most variants were rendered exclusive to gnomAD (7,646,762), followed by COSMIC (1,205,308) and ClinVar (334,517). Nevertheless, 1,720,101 SNVs were shared by at least two repositories, including 192,097 represented in all three (**Figure 2A**). These overlapping records provide opportunities to examine population, clinical and cancer-associated evidence for the same genomic alleles, while the considerably larger source-specific fractions highlight the complementary scope of the three repositories.

**Figure 2.**
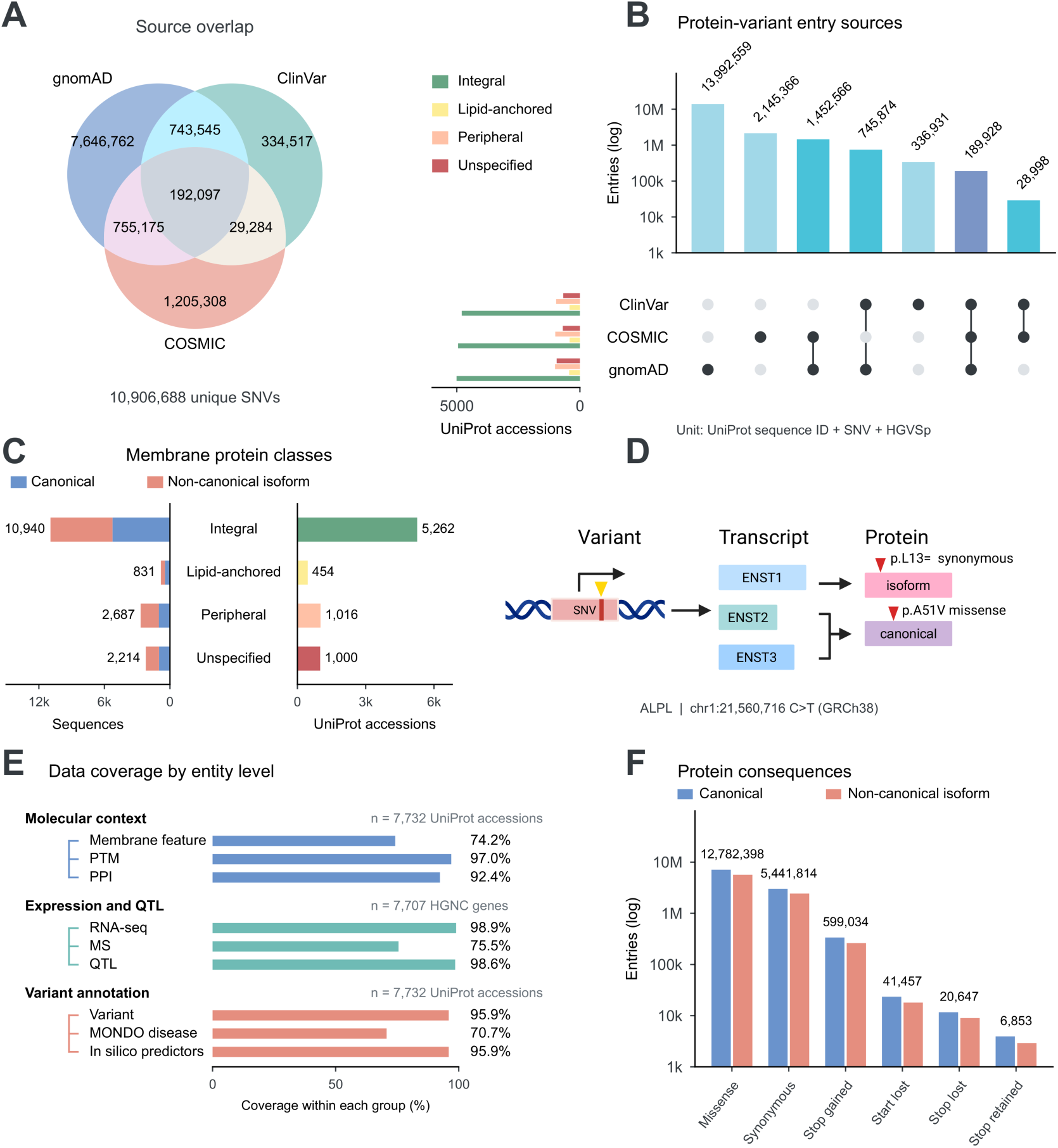
Variant sources, sequence representation and annotation coverage. (A) Overlap of 10,906,688 unique genomic single-nucleotide variants (SNVs) among ClinVar, COSMIC, and gnomAD. areas are schematic. (B) Source intersections of 18,892,222 protein-variant entries, each defined by a UniProt sequence identifier, SNV, and full HGVS protein description (HGVSp). Side bars count protein accessions by membrane class. (C) Canonical and non-canonical isoform sequences and protein accessions by membrane class. Overlapping class assignments are displayed using the priority integral, lipid-anchored, peripheral and unspecified. (D) The ALPL allele chr1:21560716 C>T has different consequences on canonical and alternative isoform sequences. (E) Annotation coverage among 7,732 protein accessions or 7,707 HGNC genes. (F) Major protein consequences in canonical and non-canonical isoform entries. Bar axes in (B) and (F) are logarithmic.

We next examined the functional consequences of coding variants using Variant Effect Predictor (VEP) annotations (**Figure 2F**). Verified consequences were retained for canonical and non-canonical isoform sequences, with each protein-variant entry defined by a UniProt sequence identifier, an SNV and the full HGVSp expression (**Figure 2D**). In total, memVar yielded 18,892,222 protein-variant entries from 10,906,688 unique genomic SNVs (**Figure 2B**). Missense variants predominated (12,782,398; 67.66%), followed by synonymous variants (5,441,814; 28.80%) and stop-gained variants (599,034; 3.17%), which introduce premature stop codons in the annotated transcripts. Less frequent consequences included start-lost variants (41,457), which may impair translation initiation, and stop-lost variants (20,647), which disrupt normal translation termination and potentially result in C-terminal protein extensions. A further 6,853 entries earned stop-retained consequences.

Beyond genomic and protein variation, memVar encompasses 7,732 membrane-associated proteins and 16,672 sequences, comprising 7,732 canonical and 8,940 non-canonical isoform sequences (**Figure 2C**). Integral membrane proteins constituted the majority (5,262), followed by 1,016 peripheral and 454 lipid-anchored proteins. The remaining 1,000 entries were classified as membrane-associated but had no specified membrane class. **T**o assess the interpretative scope of this protein collection, we further quantified annotation coverage across structural features, molecular interactions, biological functions and disease-related evidence (**Figure 2E**). Among these membrane protein in UniProt protein accessions, post-translational modification and protein-protein interaction annotations covered 97.0% and 92.4%, respectively, whereas membrane-specific features covered 74.2%. Likewise, RNA expression and QTL records covered 98.9% and 98.6% of the 7,707 HGNC genes, respectively, while mass-spectrometry-based protein abundance data covered 75.5%. Variant records and in silico predictions each covered 95.9% of protein accessions, compared with 70.7% for MONDO disease annotations, revealing a gap between the broad coverage of variant records and predictions and the availability of disease annotations. Together, these results establish the extent of variant and protein annotation within memVar while identifying specific evidence gaps that remain important for interpreting the functional and clinical consequences of membrane protein variation.

### Membrane regions and interaction sites provide molecular context

Despite the breadth of the catalogue, 820,100 of 1,299,443 ClinVar-derived SNVs (63.11%) were classified as variants of uncertain significance (VUS). This unresolved fraction highlights the gap between recording a variant and interpreting its clinical significance. For missense variants, the location of a substitution within a membrane protein constrains the molecular processes that it may affect. memVar therefore places residue annotations on the corresponding protein sequence, allowing clinical evidence to be examined alongside membrane topology, structural domains, functional sites, interaction interfaces, allele frequency, precomputed variant-effect scores, etc. Their joint presentation lays important foundations for mechanism-oriented investigation.

Topological information was derived from HTP (65), TOPDB (66), and PDBTM (67), which are three resources catalogued by UniTmp (25). Membrane depth was precomputed by OPM and lipid contacts were identified by MPLID (68). To complement the experimental evidence for the topological location of membrane proteins, we also employed DeepTMHMM2 (49) to predict transmembrane (TM) regions, re-entrant regions (RE), and interfacial helices (IF). Our data shows that 23.5% of P/LP missense substitutions and 11.1% of B/LB missense substitutions occur in UniProt-derived transmembrane segments (OR = 2.47, 95% CI, 2.36-2.59). Extending this analysis to DeepTMHMM2 predictions, we observed enrichment of P/LP relative to B/LB substitutions in TM (OR = 2.64, 95% CI = 2.52-2.76), RE (OR = 13.51, 95% CI = 10.22-18.15), and IF (OR = 2.14, 95% CI = 1.81-2.53) regions, respectively, each compared with other evaluated regions (**Figure 3B**). RE regions exhibited the strongest association, although the wider confidence interval indicates greater uncertainty in its estimated magnitude. Together, these findings reveal distinct associations between membrane topology and clinical variant classifications, highlighting membrane-associated regions that warrant closer investigation of their structural and functional constraints.

**Figure 3.**
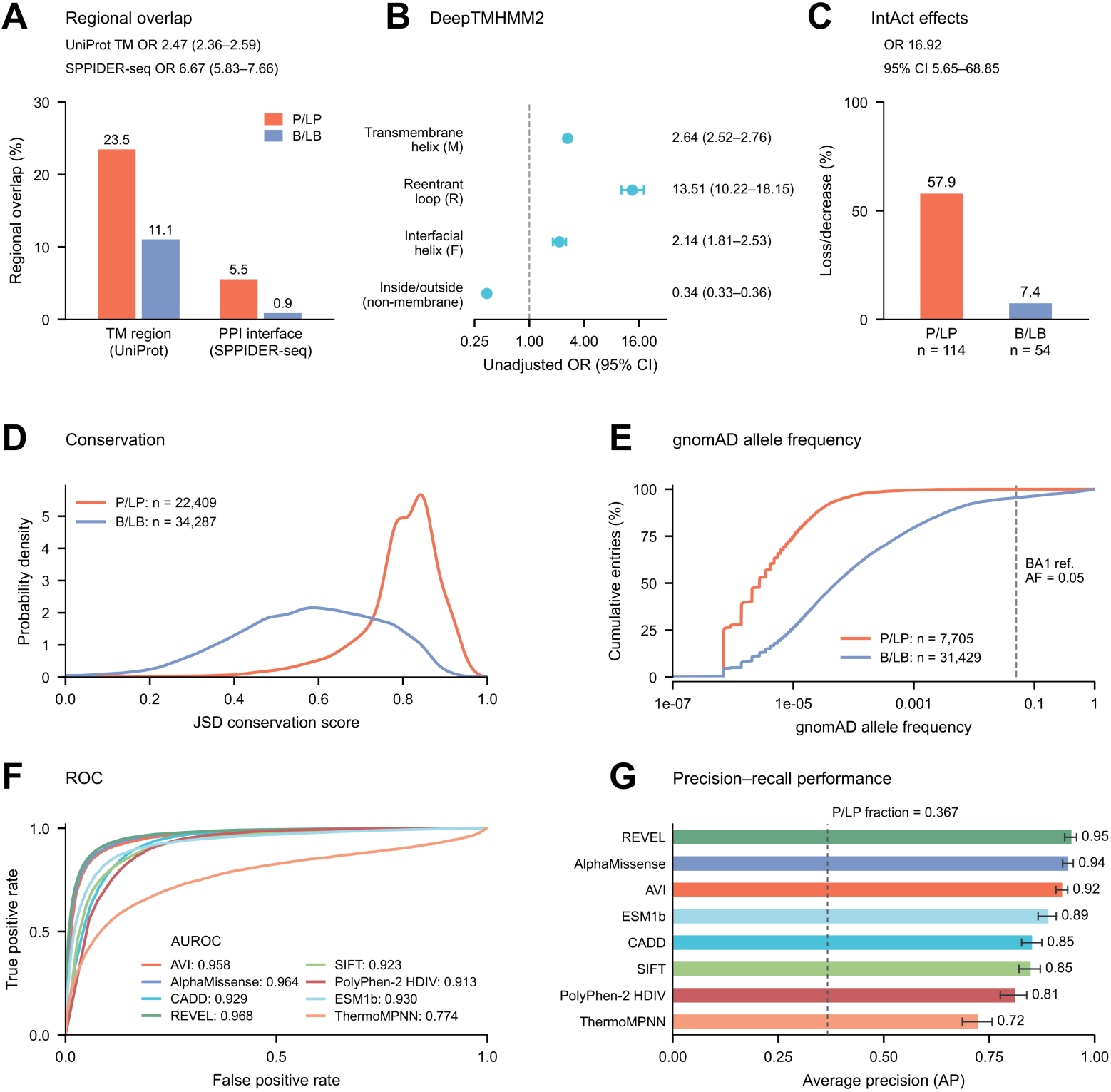
Molecular features and predictive scores of clinically classified transmembrane-protein missense variants. Pathogenic/likely pathogenic (P/LP) and benign/likely benign (B/LB) protein-variant entries are compared. (A) Overlap with UniProt transmembrane segments and SPPIDER-seq predicted protein-protein interaction (PPI) interfaces (UniProt, 21,114 P/LP and 32,011 B/LB entries. SPPIDER-seq, 21,020 and 30,657, respectively). (B) Unadjusted odds ratios (ORs) for DeepTMHMM2 regions relative to all other evaluated regions (55,708 entries). Points and error bars show ORs and 95% confidence intervals (CIs). (C) Recorded IntAct interaction loss/decrease among entries with consistent interaction-effect evidence. (D) Separately normalized Jensen-Shannon divergence (JSD) score distributions. (E) Cumulative distributions of positive gnomAD allele frequencies. The dashed line marks the 5% BA1 reference threshold (62). Counts for (C-E) are indicated in the panels. (F) Receiver operating characteristic curves and areas under the curves. (G) Precision-recall areas calculated as average precision, with 95% CIs from 1,000 bootstrap samples drawn by protein accession. Panels (F,G) use the same 40,492 entries (14,859 P/LP and 25,633 B/LB); the dashed line in (G) marks their P/LP fraction (0.367). AVI, AlphaGenome Variant Impact. CADD, Combined Annotation Dependent Depletion (92). SIFT, Sorting Intolerant From Tolerant (63). PolyPhen-2, Polymorphism Phenotyping v2 (93). HDIV, the HumDiv-trained model.

Beyond membrane topology, post-translational modifications (PTMs) provide an additional molecular context for interpreting the functional consequences of protein variation. To capture this regulatory dimension, memVar integrates PTM annotation from UniProt (39), dbPTM (69)and GlyGen (70), including 202,220 experimental records from dbPTM across 37 source-reported PTM types, such as phosphorylation, glycosylation, ubiquitination, acetylation, and methylation. PTMD 2.0 adds protein-associated PTM-disease information (71). Together, these resources enable the systematic examination of variants in relation to PTM sites and their local sequence environments, providing a basis for investigating potential disruptions to post-translational regulation and their relevance to disease (69, 72).

Protein-level annotations connect these local features to cellular function. UniProt and HPA provide localization information, and UniProt and Gene Ontology describe molecular activities and processes. Reactome annotations link these proteins to pathways involved in signal transduction, metabolism and immune system processes (73). Guide to Pharmacology target-ligand relationships and activity measurements (74), together with Rhea biochemical and transport reactions (75), place these proteins in a functional and pharmacological setting.

These annotations allow a general prediction of variant damage to be examined against specific possibilities, such as altered membrane embedding, altered modification at a PTM site or perturbation of an interaction site. The value of the integration lies in making those possibilities explicit and traceable to their supporting records.

### Pathogenic membrane protein variants are enriched at predicted interaction interfaces and associated with experimentally observed interaction defects

Protein-protein interactions (PPIs) are central to membrane protein function, and substitutions affecting interaction interfaces can compromise molecular recognition, protein complex assembly, and downstream cellular processes (76). Previous structural and functional studies have demonstrated that disease-associated missense variants are preferentially enriched at protein interaction interfaces and can selectively perturb molecular interactions without necessarily disrupting overall protein stability (35, 77). To investigate these relationships systematically across membrane proteins, memVar integrates 665,504 unique interaction pairs from BioGRID (53) and 589,765 from IntAct (54), spanning physical, association, and genetic interaction evidence. These complementary interaction networks establish the molecular partners associated with individual membrane proteins, providing a foundation for investigating how clinically classified variants relate to specific interaction interfaces.

To resolve these relationships at the amino acid level, we incorporated 310,534 partner-specific interface prediction records from SPPIDER-seq, together with structure-based interface annotations from PeSTo (56). A marked enrichment of P/LP missense substitutions emerged at predicted PPI interfaces (**Figure 3A**). Among 21,020 evaluable P/LP substitutions, 5.5% overlapped SPPIDER-seq-predicted interface residues, compared with only 0.9% of 30,657 B/LB substitutions. This corresponds to a pooled OR of 6.67 (95% CI, 5.83-7.66), indicating substantially greater interface localization among P/LP than B/LB substitutions. This pronounced difference is consistent with the functional importance of interaction surfaces, where amino acid substitutions may alter molecular recognition, binding affinity or partner specificity. Importantly, these results extend previously reported associations between pathogenic variation and protein interfaces to a membrane-protein-focused collection, including predicted interaction regions for which experimentally resolved complex structures may be unavailable.

We next investigated whether the association between clinical pathogenicity and interaction interfaces was accompanied by experimentally observed alterations in molecular interactions. Using the IMEx mutation dataset available through IntAct, we examined a separate set of substitutions with matched clinical classifications and curated experimental interaction outcomes (the rightmost panel, **Figure 3C**). Interaction loss or decrease was recorded for 57.9% of P/LP substitutions (n = 114), compared with only 7.4% of B/LB substitutions (n = 54), yielding an OR of 16.92 (95% CI, 5.65-68.85). Thus, within this experimentally characterized subset, P/LP substitutions were substantially more likely than B/LB substitutions to be associated with impaired molecular interactions. This observation accords with previous experimental studies demonstrating that disease-associated missense variants frequently perturb specific protein interactions (78), providing a potential mechanism linking individual amino acid substitutions to altered biological function (79).

Together, the pronounced enrichment of P/LP substitutions at predicted PPI interfaces and their stronger association with experimentally recorded interaction defects provide complementary evidence linking membrane protein variation to interaction-dependent functional impairment. Although the predicted-interface and experimentally characterized mutation subsets are distinct, their consistent associations with clinical pathogenicity highlight the value of integrating partner-specific interface annotations with experimentally documented interaction effects to investigate molecular mechanisms of membrane protein dysfunction.

### Disease anatomy and expression distinguish biological contexts

A molecular defect acquires disease relevance in the cells and tissues in which the affected protein operates (37). memVar connects HPA (80) and GTEx (38) RNA profiles to separate protein measurements from HPA immunohistochemistry and mass spectrometry (36). The HPA datasets used here comprise RNA profiles across 40 tissues and MS measurements across 20 tissues from one donor. GTEx (38)and QTLbase (81) additionally contribute approximately 215 million source association records spanning 21 QTL types and 7,603 genes. Tissue and study annotations identify the setting of each genetic association.

To connect recorded disease anatomy with these tissue-specific expression data, we standardized ClinVar disease conditions using MONDO and organized locations into 15 anatomical groups using UBERON (**Figure 4A**). The resulting catalogue contained 4,425 MONDO disease and related-condition concepts linked through 511,422 protein-variant entries. Within each group, we counted distinct MONDO disease concepts and protein-variant entries with P/LP, B/LB or uncertain-significance classifications (**Figure 4B**).

**Figure 4.**
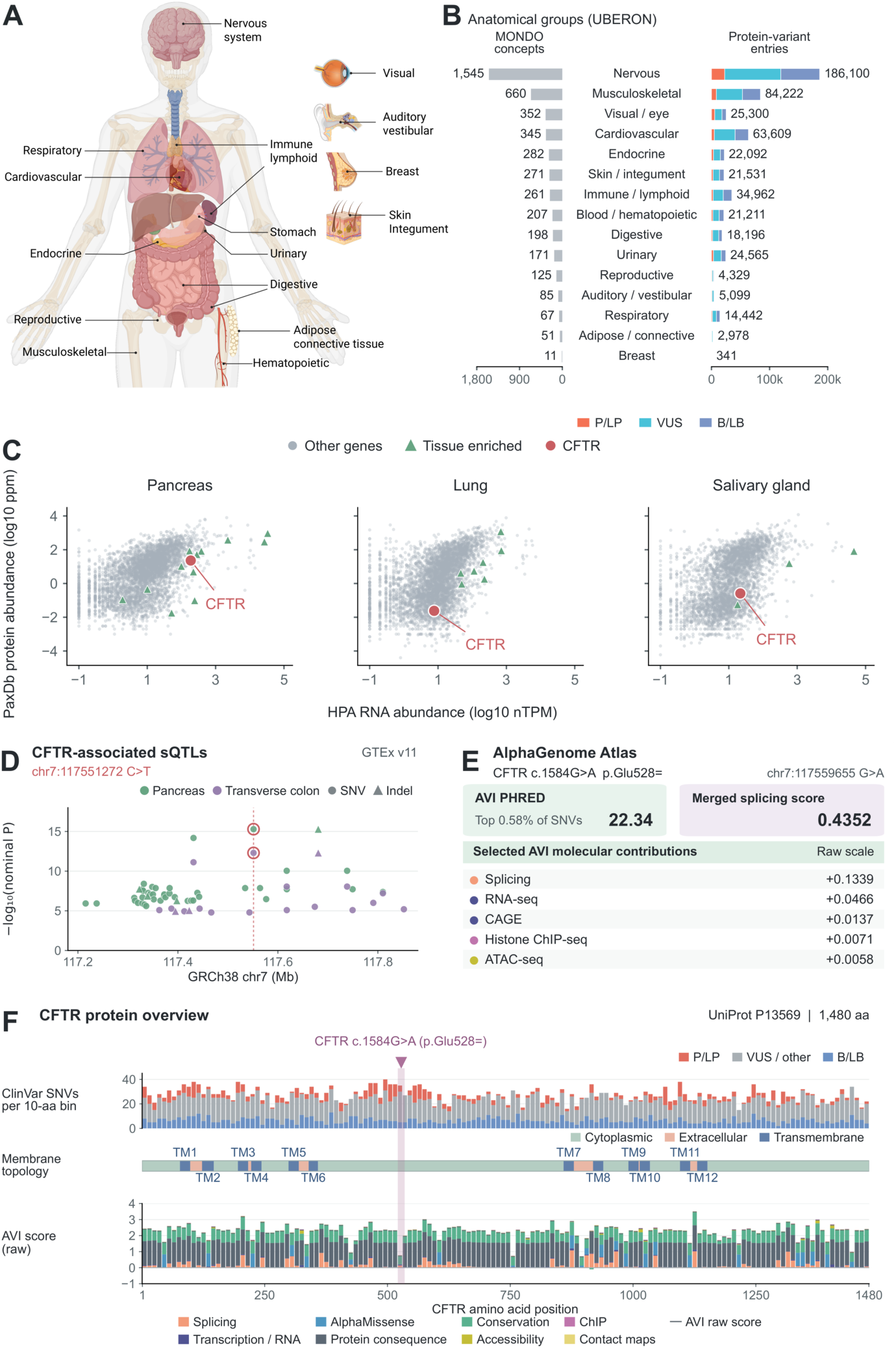
Disease, expression and regulatory contexts of membrane-protein variants. (A) Anatomical contexts used to organize disease and expression annotations. (B) Distinct MONDO concepts and clinically classified protein-variant entries across 15 broad UBERON-based anatomical groups. (C) HPA RNA abundance and PaxDb protein abundance in pancreas, lung and salivary gland, with CFTR highlighted. Triangles mark genes with ≥1 normalized transcript per million (nTPM) and ≥4-fold higher RNA abundance than in every other tissue among the 40 surveyed tissues. Protein abundance is in parts per million (ppm). (D) CFTR-associated splicing QTLs in GTEx v11. chr7:117551272 C>T is highlighted in pancreas and transverse colon. (E) Atlas AVI PHRED, merged splicing score and selected AVI feature contributions for the distinct synonymous allele CFTR c.1584G>A (p.Glu528=. chr7:117559655 G>A). (F) CFTR clinical classifications, membrane topology and AVI feature contributions along the canonical sequence. Clinical counts use 10-residue bins. AVI tracks show one representative variant per bin, selected by highest raw AVI score except for the highlighted p.Glu528= bin. Genomic positions use GRCh38. CAGE, cap analysis of gene expression. ChIP-seq, chromatin immunoprecipitation sequencing. ATAC-seq, assay for transposase-accessible chromatin sequencing.

The resulting distributions describe recorded associations rather than disease prevalence or organ-specific risk. Within the P/LP, B/LB and VUS subset, the nervous-system group contained 1,545 MONDO concepts and 186,100 protein-variant entries, followed by the musculoskeletal group with 660 concepts and 84,222 entries (**Figure 4B**). This shared anatomical organization provides a route from a variant-associated condition to relevant expression backgrounds.

CFTR provides a concrete example (82). Its variant-linked disease conditions in memVar include cystic fibrosis (MONDO:0009061) and hereditary chronic pancreatitis (MONDO:0008185). HPA RNA profiles and PaxDb protein-abundance measurements showed higher CFTR abundance in pancreas than in lung or salivary gland (**Figure 4C**). RNA and protein abundance were compared separately on their respective measurement scales. These profiles provide tissue-expression context for investigating CFTR variant effects and selecting relevant experimental systems.

It has been suggested that transcription, RNA processing, translation and protein turnover can all contribute to differences between RNA and protein abundance (83, 84). The present tissue comparisons do not isolate these mechanisms. QTL annotations therefore supply a separate genetic perspective on expression, splicing and other molecular traits, with tissue and study information retained so that an association can be evaluated in its original context (85).

A local CFTR example highlights chr7:117551272 C>T, an sQTL associated with the same splice-junction phenotype in pancreas and transverse colon, with nominal *P* = 5.13 × 10⁻¹⁶ and 5.01 × 10^−13^ (**Figure 4D**). These tissue-specific associations identify a transcript-processing phenotype for functional investigation, while further evidence is needed to establish whether the allele alters splicing and whether the associated splice change contributes to disease. Documented gene-disease evidence is available alongside these exploratory annotations. OMIM, MedGen, ClinGen, GenCC, MONDO and HPO connect gene-disease relationships, inheritance, curation strength and clinical phenotypes through explicit identifiers. ClinGen dosage-sensitivity records remain separate, and ClinGen assertions distributed through GenCC retain their provenance rather than being counted as independent evidence. For CFTR, the retained ClinGen curation describes a definitive association with autosomal recessive cystic fibrosis and a loss-of-function mechanism. This provides established disease context against which tissue and regulatory hypotheses can be considered.

We further added the information about regulatory predictions around the genes encoding membrane proteins. We incorporated variant-impact and splicing predictions from AlphaGenome Atlas. AlphaGenome Atlas AVI scores map to 10,863,110 of the 10,866,094 SNVs (99.97%), accompanied by 18 source feature contributions. Merged splicing scores cover 10,762,279 SNVs (45). AVI incorporates features including predicted splicing and chromatin effects, AlphaMissense scores and conservation, so agreement between AVI and its component predictors is not independent evidence. A second CFTR allele, c.1584G>A (p.Glu528=), illustrates how these predictions can inform the investigation of a synonymous variant (**Figure 4E, F**). Its Atlas splicing score is 0.4352 while AVI PHRED score is 22.34, with a positive contribution from the splicing feature and substantial contributions from genomic conservation (**Figure 4E**). The CFTR protein view displays this allele, which has conflicting ClinVar classifications, alongside membrane topology and AVI feature contributions (**Figure 4F**). Although the encoded amino acid is unchanged, the splicing prediction motivates functional investigation of possible effects on RNA processing.

### CFTR variants illustrate mechanism-oriented exploration

To illustrate how memVar supports mechanism-oriented variant interpretation, we examined two CFTR substitutions with contrasting clinical annotations, including the pathogenic gating variant G551D and the variant of uncertain significance W202R. Querying CFTR (UniProt accession code: P13569) retrieves a protein record that integrates molecular, pathogenic, and clinical annotations at the sequence, topological, structural levels in multiple visualization modes (**Figure 5**). These complementary views allow users to contextualize an established disease mechanism and formulate hypotheses for a variant whose consequences remain unresolved functionally.

**Figure 5.**
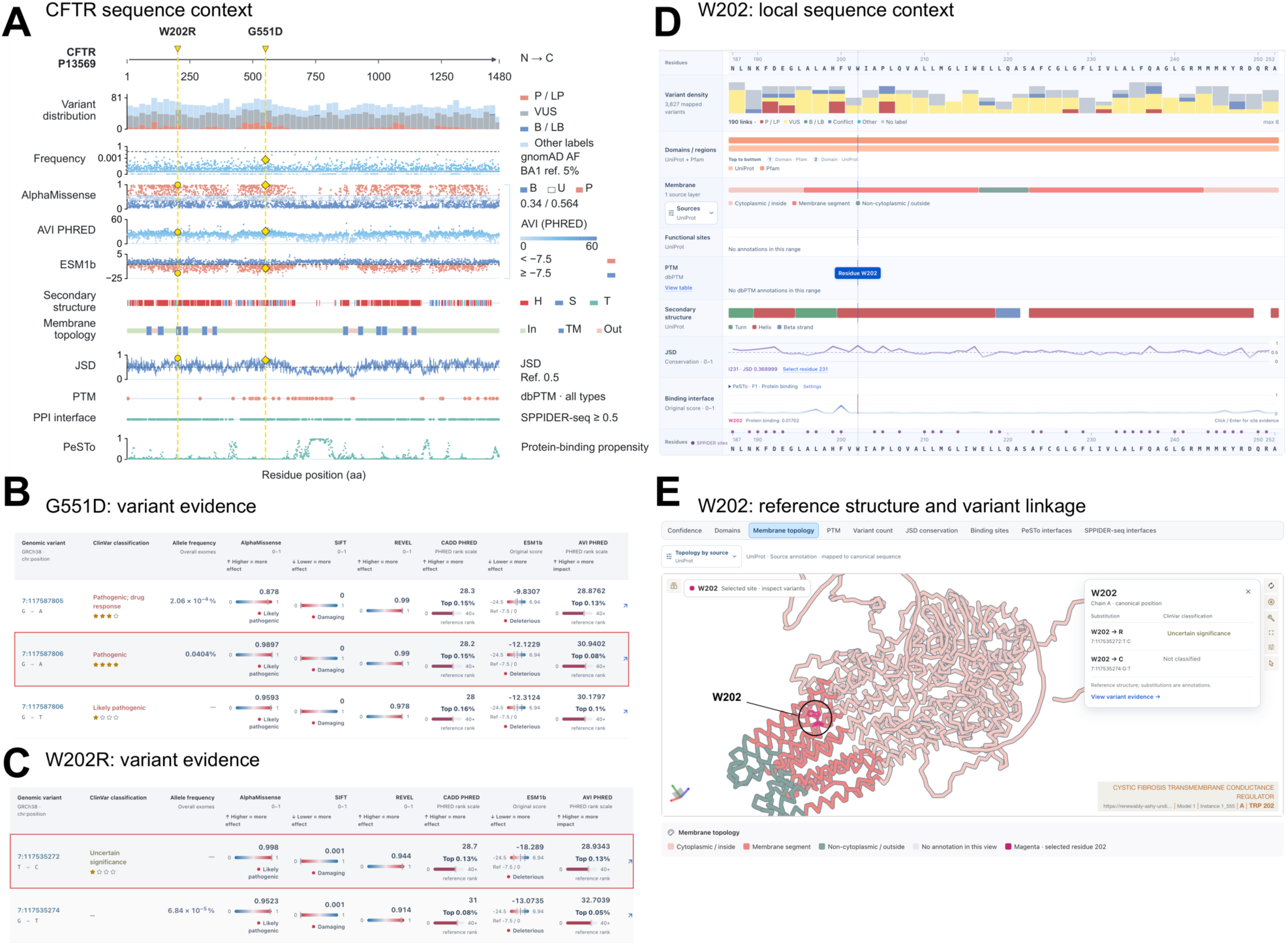
Linked sequence, variant and structure views illustrated by CFTR. (A) Sequence overview combining variant distributions, allele frequencies, predictive scores, secondary structure, membrane topology, conservation, PTMs and interface predictions. (B,C) Variant evidence for G551D and W202R, respectively. (D) Local sequence annotations around W202. (E) W202 highlighted in the reference structure, with links to substitutions at that residue. The structure displays the reference sequence. Clinical labels and scores are those shown in the interface snapshots. CADD, Combined Annotation Dependent Depletion (92). SIFT, Sorting Intolerant From Tolerant (63). REVEL, Rare Exome Variant Ensemble Learner (64).

For G551D, we examined how various annotations are integrated to assist in fathom its experimentally established impairment of ATP-dependent gating (10). In the sequence view of memVar, G551 is rendered in the nucleotide-binding domain 1, which is outside the transmembrane segments (**Figure 5A**). G551D is clinically classified as pathogenic by ClinVar, in consistency with deleterious predictions by AlphaMissense (0.990), REVEL (0.990), and AVI PHRED (30.94) (**Figure 5B**). Using ThermoMPNN, a destabilizing change (ΔΔG) of +0.79 kcal/mol is additionally predicted. The meta-analysis of the variant effect of G551D yields plausible explanation for the impaired ATP-dependent gating.

We then examined W202R, a mutation that occurs at position 202 within transmembrane helix 3 (residues 196-216). Its related information is shown across multiple panels in a calibrated sequence, such as the PesTo-predicted interfacial potential (**Figure 5D**). Our precomputed Jensen-Shannon divergence (JSD) score of 0.873 implies strong evolutionary conservation at this position. Substitution of this conserved aromatic residue with a positively charged arginine suggests altered local packing or membrane-protein biogenesis as testable hypotheses. In addition, by interpreting these prediction scores in their structural context, we can formulate testable hypotheses about local packing and protein biogenesis (**Figure 5E**). While W202R is grouped as VUS, it is suggested as being pathogenic by predicted variant effects, 0.998, 0.944, −18.29, and 28.93 for AlphaMissense, REVEL, ESM1b, and AVI PHRED, respectively (**Figure 5C**), together with a ThermoMPNN-predicted ΔΔG of +2.34 kcal/mol. Incorporation of these predictions with the transmembrane location supports prioritizing W202R for measurements of CFTR maturation, cell-surface abundance and channel activity. These complementary readouts could help distinguish impaired biogenesis or trafficking from functional defects related to membrane proteins.

Altogether, the two examples illustrate how effects of variants from clinical annotations are interpreted in molecular contexts and under functional hypotheses. Users can further consult expression, regulatory, interaction, and disease annotations in support of their experimental choices and schemes, while retaining the source clinical classification throughout the analysis.

## Discussion

memVar brings existing variant observations into a common molecular and disease framework for human membrane-associated proteins. Its contribution is the coordination of evidence that is usually consulted separately: an allele and its source assertions, the affected residue and its membrane environment, and the interactions, tissues and diseases associated with the protein. Earlier resources established the value of membrane-specific curation (19–21), and general platforms have made variant-to-protein mapping increasingly accessible (26, 27). memVar extends this organization across a broad membrane-associated reference set and links membrane features to interaction, expression and regulatory context. The CFTR examples show how these connections can turn a general prediction of damage into a more specific question about protein biogenesis or function.

The current analyses characterize the integrated resource rather than validate a new clinical predictor. Enrichment of P/LP substitutions in membrane segments and predicted interfaces is consistent with prior work, but the pooled comparisons remain sensitive to gene composition, clinical ascertainment and annotation coverage. Cluster bootstrap intervals account for the specified dependence structure without removing these biases. Source clinical classifications can change, and the present comparisons do not impose a further ClinVar review-status filter. Missing membrane annotations are also distinct from evidence that a feature is absent. These constraints should guide interpretation of both the aggregate results and individual variant records.

Direct functional evidence remains the most important area for expansion. We plan to incorporate suitable multiplexed assays of variant effect, including data available through ClinMAVE (86), and to extend literature curation of variant-associated phenotypes and mechanisms. For membrane proteins, assays of maturation and trafficking, transport or channel activity, and signalling can distinguish consequences that generic pathogenicity scores do not resolve (87). Experimental system, assay endpoint and score definition must remain attached to each measurement because an assay phenotype is not equivalent to a clinical diagnosis (88). The same discipline applies to tissue and regulatory data: bulk-tissue expression, QTL associations and reference-sequence predictions provide complementary context but do not by themselves identify the causal allele, cell type or molecular pathway. We also plan to extend topology, membrane-interface and lipid-contact coverage while retaining source confidence and mapping information.

A second priority is independent evaluation of computational evidence in defined membrane contexts. EVE, ESM1b, and AlphaMissense capture evolutionary or structural constraints, while membrane-specialized methods incorporate additional topological or sequence information (32, 89). Existing AlphaMissense assessments already demonstrate the importance of reference-data quality (30). Future benchmarks should separate related proteins between development and test sets, address overlap with predictor training data, and use independently measured functional effects where possible (90). Performance and calibration should be assessed by protein class, membrane region and experimental endpoint, with annotation missingness reported explicitly. Agreement between predictors, particularly those sharing features or training data, cannot substitute for validation (91). These datasets will also support investigation of interpretable scoring approaches that combine local membrane features with functional and disease evidence. By retaining these evidence distinctions, memVar provides a framework in which new measurements can accumulate, unresolved mechanisms can be identified, and membrane protein variants can be reassessed as knowledge develops.

## Acknowledgements

We thank the High-Performance Computing (HPC) platform at The Hong Kong University of Science and Technology (Guangzhou) for providing computational resources.

## Declarations

The authors declare no competing interests.

## Data availability

The memVar website is accessible at https://memvar.sunnylab.org.

## Code availability

The code used to develop memVar is made publicly available at https://github.com/sunnylab-cb/memvar.

## Author contributions

J.S. designed this study. Y.Z. investigated the literature, curated the data, performed the data analysis, and prepared the draft. Y.Z. developed the memVar website with input from S.H. and J.Y. ColabFold structures deposited in memVar were generated by J.Y. and S.H. J.S. provided supervision, investigated the literature, interpret the analysis, and revised the manuscript. All authors have given approval to the final version of the manuscript.

