## Supplementary Material for "memVar: an integrated resource for interpreting human membrane protein variation in molecular and disease contexts"

**Supplementary Method 1**

We collected canonical and isoform sequences from reviewed human UniProtKB/Swiss-Prot entries with the Membrane keyword (KW-0472). This selection included peripheral and lipid-anchored proteins as well as transmembrane proteins. Single-nucleotide variants (SNVs) were collected from ClinVar, COSMIC and gnomAD exomes and normalized against GRCh38 with bcftools 1.24 (Supplementary Table S1). We retained only SNV annotations with a protein-level consequence.

Source records differed in both transcript reference sets and versions, with the ClinVar and COSMIC records used here referencing RefSeq and Ensembl transcripts, respectively. To normalize variant annotations across sources, we used Ensembl Variant Effect Predictor (VEP) 116.0 to reannotate variants for all transcripts supported by each source record, while retaining the original annotations. We matched Ensembl transcripts by stable identifier and RefSeq transcripts through MANE Select or Plus Clinical pairs. An exact match to the full UniProt sequence was required, and we checked the residues or boundaries in the reported protein change. Either the original annotation or its VEP counterpart could meet these criteria. Each memVar entry represents one UniProt sequence, one SNV and one HGVS protein description (HGVSp).

**Supplementary Method 2**

gnomAD allele frequencies, dbSNP identifiers and AlphaGenome Atlas scores were matched by GRCh38 position and alleles. Atlas provides the AlphaGenome Variant Impact (AVI) score, its feature contributions and a splicing score. dbNSFP scores were matched by allele. Transcript-specific scores also required the same transcript, protein and amino acid change (Supplementary Table S5). Conflicting or unavailable scores remained missing.

**Supplementary Method 3**

We mapped membrane regions and post-translational modifications (PTMs) by UniProt residue position. For structural annotations, we used SIFTS to map residues and checked their identity against UniProt. Pfam domains were identified with HMMER using the gathering threshold for each model, and membrane topology was predicted with DeepTMHMM2 (Supplementary Tables S2 and S4).

For each protein sequence, we searched UniRef90 with MMseqs2, requiring an E-value $\leq 10^{-5}$ and $\geq$60% query coverage. After excluding human representatives and exact query matches, we clustered homologues at 95% identity and aligned up to 150 representatives to the query using MAFFT E-INS-i. Jensen–Shannon divergence (JSD) and Zvelebil conservation scores were calculated using only the homologous sequences. Positions with no standard amino acid among the homologues remained unscored.

**Supplementary Method 4**

Functional annotations were linked by gene or protein identifier and IntAct mutation effects by protein residue. SPPIDER-seq provided residue-level interface probabilities for protein pairs and PeSTo estimated binding propensities at individual residues. We computed ThermoMPNN stability estimates by canonical sequence, structure or fragment, and amino acid substitution.

AlphaFold DB structures were matched by sequence. We predicted a structure for each membrane protein using ColabFold. Local MMseqs2 searches against UniRef30 (February 2023) and the ColabFold environmental sequence database (August 2021) provided unpaired alignments. For each sequence, we ran AlphaFold2-pTM with one model, one random seed (0) and three recycles, without templates or ensembling. Predictions were run on an NVIDIA A100 GPU with TensorFloat-32 enabled. pLDDT was used for ranking, and Amber relaxation was omitted.

**Supplementary Method 5**

To standardize disease and phenotype conditions, we mapped MONDO identifiers from ClinVar to current disease terms, accepting alternative identifiers and obsolete terms with a single replacement. Each ClinVar classification remained associated with the reported condition or condition set.

We used anatomical locations annotated to each MONDO term or inherited from its ancestors. For each location, we followed UBERON subclass and part-of relationships to the most specific applicable groups among the 15 defined for memVar (Supplementary Table S6). A disease could be assigned to more than one anatomical group. Tissues from the Human Protein Atlas (HPA), GTEx RNA and PaxDb were assigned to the same groups where an anatomical match was available. Quantitative trait loci (QTLs) from GTEx and QTLbase were linked by target gene, retaining the original tissue labels and genome assemblies (Supplementary Table S3).

**Supplementary Method 6**

We restricted the analyses in Figure 3 to ClinVar-classified missense entries in proteins with the UniProt Transmembrane keyword (KW-0812). Of these, 22,409 were classified as pathogenic or likely pathogenic (P/LP) and 34,292 as benign or likely benign (B/LB). Each analysis used entries with the relevant annotation. A residue was classified as an interface site if SPPIDER-seq assigned a probability ≥0.5 for at least one partner with the query protein as receptor. DeepTMHMM2 predicts residue-level membrane topology from protein sequences. After excluding signal peptides, transit peptides and transmembrane β-strands, we compared transmembrane helices (TM), re-entrant regions (RE), interfacial helices (IF) and non-membrane regions (inside/outside), testing each category against the other three combined. The IMEx mutations dataset available through IntAct records experimentally observed effects of sequence changes on protein interactions. For this analysis, reports on the same substitution had to agree on loss or weakening of an interaction, or on no effect.

We calculated conditional exact odds ratios with 95% confidence intervals and used two-sided Fisher exact tests. Bonferroni correction was applied separately to the three regional tests and the four DeepTMHMM2 comparisons. JSD density estimates used a Gaussian kernel (bandwidth 0.02) with reflection at 0 and 1. Allele frequency curves included only positive gnomAD exome frequencies for the full exome cohort.

We compared eight predictors (SIFT, PolyPhen-2 HDIV, CADD, REVEL, ESM1b, AlphaMissense, AlphaGenome Atlas AVI and ThermoMPNN) on the same 40,492 entries with all eight scores available. SIFT and ESM1b scores were reversed so that higher values indicated greater predicted impact. Predictor performance was evaluated using the area under the receiver operating characteristic curve and average precision. Figure 3G shows 95% confidence intervals for average precision from 1,000 paired bootstrap samples of UniProt accessions.

For Figure 4B, we counted distinct MONDO concepts and entries classified as P/LP, B/LB or uncertain significance within each anatomical group. Each unique combination of primary UniProt accession, SNV and full HGVSp was counted once per group. Figure 4C pairs HPA RNA and PaxDb protein abundance within a tissue where both values were positive and the UniProt match was unambiguous. Among the 40 HPA tissues, the tissue with the highest RNA abundance was marked if this value was ≥1 nTPM and at least four times the second-highest value, following the HPA fourfold criterion for "tissue enriched" expression.

Figure 4D uses significant CFTR splicing QTL pairs reported by GTEx. In Figure 4F, we counted 4,037 SNVs with ClinVar records in 10-residue bins along canonical CFTR. Among all 5,560 mapped SNVs, we selected the one with the highest raw AVI score in each bin. For residues 521 to 530, we show the example variant c.1584G>A (p.Glu528=).

**Supplementary Table S1**. Reference and variant datasets incorporated into memVar.

| Resource | Version or date | Data included |
|---|---|---|
| UniProtKB/Swiss-Prot | 2026_03 | 7,732 accessions with 7,732 canonical and 8,940 non-canonical sequences, plus sequence features, topology and cross-references |
| HGNC | 13 September 2026 | Gene names and 7,707 HGNC identifiers linked to memVar proteins |

| Resource | Version or date | Data included |
| --- | --- | --- |
| Ensembl and VEP | 116 and VEP 116.0 | GRCh38 reference models and 23,292,622 verified associations between SNVs and transcripts |
| MANE | 1.5 | RefSeq–Ensembl Select and Plus Clinical transcript pairs |
| ClinVar variant summary | 6 September 2026 | 1,299,443 SNVs with clinical annotations |
| ClinVar condition records | 5 September 2026 | 1,747,134 records linking variants to reported conditions or condition sets |
| COSMIC | 104 | 4,551,304 annotations for 2,181,864 GRCh38 SNVs, with tumour site and classification |
| gnomAD exomes | 4.1.1 | 29,066,408 annotations for 9,337,579 SNVs, plus frequencies for 9,337,876 alleles |
| dbSNP | Build 157 | rs identifiers for GRCh38 SNVs |
| dbNSFP | 5.4a | 18 predictors across 34,199,310 records for individual SNVs or SNV-transcript pairs |
| AlphaGenome Atlas | September 2026 | AVI scores for 10,905,655 SNVs and splicing scores for 10,812,739 SNVs, with available contribution scores |

Note. The three variant sources overlap. Together they account for 10,906,688 distinct SNVs and 18,892,222 protein-variant entries. We matched gnomAD frequencies by allele independently of the reported protein consequence. The dbNSFP total includes records with missing scores.

**Supplementary Table S2**. Molecular, interaction and disease resources incorporated into memVar.

| Resource | Version or date | Data included |
| --- | --- | --- |
| TOPDB | 3.3 | 202,395 topology-region records |
| HTP | d.2.2 | 807,796 topology-region records |
| PDBTM | 20250404 | 262,614 membrane-region records |
| TmAlphaFold | 1.1.1 | 870,888 CCTOP regions, 62,237 TMDET regions, 5,049 model evaluations |
| TOPDOM | 3.3 | 9,020 topology-domain records |
| OPM | | 2,935,154 residue observations in structures |
| MPLID | 2.0 | 1,148 structure records and 1,264,315 residue records with contact status |
| BioDolphin | vr1.1 | 16,287 complex/interaction records and 49,445 site records |
| AlphaFold DB | v6 | 8,824 sequence-matched models, comprising 7,530 full-length and 1,294 fragment matches |
| SIFTS | | Residue mapping between structures and UniProt sequences |
| dbPTM | 2025 | 202,220 experimental PTM records across 37 PTM labels |
| PTMD2 | 2.0 | 100,468 records linking PTMs to disease across 54 PTM labels |
| GlyGen | 2.11.1 | 38,612 glycosylation records |
| GO and GOA | 26 July 2026 (GO), 28 July 2026 (GOA) | 597,211 annotations, with ontology terms and relationships |

| Resource | Version or date | Data included |
|---|---|---|
| Reactome | 97 | 80,300 pathway associations, with pathway hierarchy and descriptions |
| Rhea | 142 | 13,145 protein–reaction relationships, with reaction and compound annotations |
| GtoPdb | 2026.2 | 15,516 interactions, with target and ligand annotations |
| BioGRID | 5.0.262 | 665,504 records from the full download and 662,140 from the interaction context collection |
| IntAct / IMEx | 9 January 2026 | 589,765 records from the full download, 169,835 interaction context records and 28,833 mutation effect records |
| MONDO | 6 July 2026 | 4,425 disease and related-condition concepts, 742,926 ClinVar–MONDO links |
| UBERON | 19 June 2026 | Anatomical ontology relationships used to group diseases and tissues |
| OMIM | 14 July 2026 | 2,532 genemap2, 3,252 mim2gene, 3,252 mimTitles and 3,658 morbidmap records |
| KEGG DISEASE | 28 September 2026 | 1,481 disease records with identifiers and reference links |
| HPO | 23 June 2026 | 102,266 phenotype annotations |
| GenCC | 27 July 2026 | 16,300 gene–disease submissions |
| ClinGen | 13 to 14 July 2026 | 2,359 gene–disease, 2,909 activity-summary and 663 gene-curation records |
| MedGen | 13 July 2026 | 23,998 identifier mappings, 5,724 names, 4,268 definitions and 2,668 merged-identifier records |

Note. UniProt also contributed 195,821 sequence feature records, 41,968 topology records and 46,379 PTM features or comments. After cross-database deduplication, structured records from UniProt, dbPTM, PTMD2 and GlyGen covered 72 PTM types, with each covalent modification family counted as one type. Within BioGRID and within IntAct, the interaction collections overlap.

**Supplementary Table S3**. Expression and regulatory datasets incorporated into memVar.

HPA release 25.1 uses Ensembl 109 gene annotations. All GTEx datasets are from v11.

| Dataset | Source | Measurement | Data included |
|---|---|---|---|
| Normal-tissue RNA | HPA | nTPM by gene and tissue | 302,040 records across 40 tissues |
| FANTOM5 tissue expression | HPA | CAGE normalized tags per million | 312,340 records across 46 tissues |
| Single-cell-type RNA | HPA | nCPM by gene and cell type | 1,158,234 records across 154 cell types |
| Cancer RNA | HPA | pTPM by gene and sample | 62,562,291 records across 31 cohorts |
| Normal-tissue immunohistochemistry | HPA | Staining by antibody, tissue and cell type | 441,880 records |
| Cancer immunohistochemistry | HPA | Staining categories and patient counts | 110,819 records across 20 cancer types |

| Dataset | Source | Measurement | Data included |
|---|---|---|---|
| Normal-tissue mass spectrometry | HPA | Protein intensity by tissue and replicate | 445,320 records across 20 tissues |
| Deep Visual Proteomics | HPA | Protein intensity by cell type | 200,394 records across 27 contexts |
| CPTAC cancer proteomics | HPA | Protein $\log_2$ fold change and adjusted P-value by cancer type | 37,459 records across 11 cancer types |
| Tissue RNA | GTEx | Median TPM by gene and tissue | 7,601 genes across 68 tissue columns |
| Integrated protein abundance | PaxDb 6.1 | Protein abundance in parts per million | 172,439 records across 52 integrated datasets |
| eQTL | GTEx | Variant–gene association | 23,608,070 associations |
| sQTL | GTEx | Variant–splice-junction association | 34,306,280 associations |
| apaQTL | GTEx | Variant–polyadenylation association | 4,835,546 associations |
| Molecular QTLs | QTLbase v2 | Variant–molecular-target association | 151,910,444 associations across 21 QTL types |

Note. The 214,660,340 QTL associations span 50 tissue contexts in GTEx and 84 contexts in QTLbase. QTLbase includes metadata for 1,681 studies. Expression record counts include rows with missing values. HPA reports normalized transcripts per million (nTPM), normalized counts per million (nCPM) and protein-coding transcripts per million (pTPM).

**Supplementary Table S4**. Computed molecular annotations included in memVar.

| Annotation | Method or reference | Data included |
|---|---|---|
| Protein domains | HMMER 3.4, Pfam 38.2 | 53,571 hits from 3,688 Pfam models |
| Membrane topology | DeepTMHMM2 | 114,337 segments across 16,672 sequences |
| Residue conservation | Capra JSD, AACon 1.1 and UniRef90 2026_02 | 9,823,117 residue positions, including positions without conservation scores |
| Partner-specific interfaces | SPPIDER-seq | Interface predictions for 310,534 protein sequence pairs |
| Binding propensities | PeSTo, AlphaFold DB v6 | Binding propensity predictions for 5,792,650 residues |
| Stability changes | ThermoMPNN default model, AlphaFold DB v6 | Predicted stability changes for 6,292,698 amino acid substitutions |

**Supplementary Table S5**. Variant-effect prediction scores obtained from dbNSFP 5.4a.

Score denotes the predictor's numerical output, with scales and directions that vary by method. Rank denotes the relative position of a score among those from the same predictor in dbNSFP, expressed on a 0–1 scale with higher values indicating greater predicted deleteriousness. Rank scores apply to the SNV and are not probabilities of pathogenicity. Classification denotes the categorical prediction recorded in dbNSFP, such as damaging or tolerated for SIFT. MisFit reports two scores, D and S. PolyPhen-2 HDIV and HVAR are separate models.

| Predictor | Raw score level | Data included |
|---|---|---|
| SIFT | Transcript | Score, rank, classification |
| PROVEAN | Transcript | Score, rank, classification |

| Predictor | Raw score level | Data included |
|---|---|---|
| PolyPhen-2 HDIV | Transcript | Score, rank, classification |
| PolyPhen-2 HVAR | Transcript | Score, rank, classification |
| CADD | SNV | Raw score, PHRED score, rank |
| MetaRNN | Transcript | Score, rank, classification |
| REVEL | Transcript | Score, rank |
| MutPred2 | Transcript | Score, rank, classification, five highest-ranked mechanisms |
| gMVP | Transcript | Score, rank |
| MPC | Transcript | Score, rank |
| ALoFT | Transcript | Tolerant, recessive and dominant probabilities, classification, confidence and transcript proportions |
| ClinPred | SNV | Score, rank, classification |
| ESM1b | Transcript | Score, rank, classification |
| AlphaMissense | Transcript | Score, rank, classification |
| MutFormer | SNV | Score, rank |
| MisFit | Transcript | D and S scores and ranks, permissive and stringent D classifications |
| popEVE | Transcript | Score, rank, classification |
| GPN-MSA | SNV | Score, rank, classification |

**Supplementary Table S6**. UBERON terms used to define the 15 anatomical groups.

UBERON provides standardized terms and identifiers for anatomical structures. The terms listed in each row are used to define the corresponding anatomical group in memVar (Supplementary Method 5). The UBERON prefix is omitted from the table. For example, 0001016 (nervous system) and 0000955 (brain) are both included in the nervous system group.

| Anatomical group | UBERON identifiers |
|---|---|
| Nervous system | 0001016, 0000955, 0002240, 0000010 |
| Visual system | 0002104, 0000970, 0004088 |
| Auditory and vestibular system | 0016490, 0004681, 0001690 |
| Cardiovascular system | 0004535, 0015410, 0000948, 0001133, 0001981, 0002049, 0007798 |
| Respiratory system | 0001004, 0002048, 0000004 |
| Digestive system | 0001007, 0005409, 0000165, 0001723, 0001264 |
| Endocrine system | 0000949, 0000007, 0002369, 0001132, 0002046 |
| Urinary system | 0001008, 0002113, 0001255, 0000056, 0000057 |
| Reproductive system | 0000990, 0001987 |
| Blood and haematopoietic system | 0002390, 0000178, 0002371 |
| Immune and lymphoid system | 0002405, 0002465, 0006558, 0000029, 0002106, 0002370, 0002372 |
| Musculoskeletal system | 0002204, 0001434, 0001134, 0001135, 0002385, 0002481, 0002418, 0014892 |
| Skin and appendages | 0002416, 0002097 |
| Breast | 0000310 |
| Adipose and connective tissues | 0001013, 0002384 |
